# Infection with non-retroviral RNA viruses produces virus-derived DNA across diverse insect species

**DOI:** 10.64898/2026.03.30.715342

**Authors:** Anton Bilsen, Sara Ottati, Cecilia Parise, Simona Abbà, Jozef Vanden Broeck, Luciana Galetto, Dulce Santos

## Abstract

Growing evidence shows that eukaryotic genomes contain DNA sequences of non-retroviral RNA virus origin, yet the mechanisms underlying the generation of this virus-derived complementary DNA (vDNA) remain poorly understood, particularly in insects. Here, we demonstrate that infection with diverse non-retroviral RNA viruses induces the production of reverse-transcribed vDNA across multiple lepidopteran and hemipteran species. We further show that a *Drosophila melanogaster*–derived cell line actively secretes reverse-transcribed vDNA associated with extracellular vesicles (EVs), widely conserved nanoscale mediators of intercellular communication. Together, these findings reveal that infection-induced vDNA formation is a widespread phenomenon in insects and suggest that EVs may facilitate the systemic dissemination of vDNA.

## Introduction

Reverse transcription (*i.e.*, the flow of genetic information from RNA to DNA) is a hallmark of retroviruses and retrotransposons. These genetic mobile elements encode RNA-dependent DNA polymerases, named reverse transcriptases, that catalyze the synthesis of DNA starting from an RNA template as part of their replication strategy (Telesnitsky & Goff, 1997).

With the advance of genomic technologies, virus-derived complementary DNA (vDNA) has been recurrently identified in eukaryotes, including sequences derived from non-retroviral RNA viruses with double- and single-stranded (positive and negative sense) genomes. This DNA is generally found in the form of endogenous viral elements (EVEs), which consist of partial fragments of vDNA integrated in the host genome (Holmes, 2011). Since non-retroviral RNA viruses do not encode reverse transcriptases, thereby not going through a DNA stage during their cycle, the circumstances and mechanisms leading to their vDNA (and potentially EVE) formation remain unclear. However, studies in various dipteran insects suggest that endogenous reverse transcriptases in the host may mediate vDNA formation (Goic et al., 2013; Goic et al., 2016; Tassetto, Kunitomi, & Andino, 2017; Tassetto et al., 2019).

EVEs from non-retroviral viruses have been identified in every insect genome for which they have been investigated (*e.g.* Blair, Olson, & Bonizzoni, 2019; Chen et al., 2015; Crochu et al., 2004; Flynn & Moreau, 2019; Lequime & Lambrechts, 2017; Maori, Tanne, & Sela, 2007; Palatini et al., 2017; Suzuki et al., 2017; ter Horst et al., 2019). Nevertheless, the event of vDNA generation upon infection with non-retroviral RNA viruses has been experimentally observed only a limited number of times. Specifically, this has been reported in fruit flies and mosquitoes, in cell lines derived from these species (Goic et al., 2013; Goic et al., 2016; Mondotte et al., 2018; Mondotte et al., 2020; Nag, Brecher, & Kramer, 2016; Poirier et al., 2018; Tassetto, Kunitomi, & Andino, 2017; Tassetto et al., 2019), and in a silkworm (*Bombyx mori*)-derived cell line (Zhu et al., 2022). While such findings have been explored to some extent in these species, the degree to which vDNA generation is conserved across insect taxa remains unresolved.

In addition, considering a potential role of vDNA as a modulator of antiviral immunity (Goic et al., 2013; Goic et al., 2016; Tassetto, Kunitomi, & Andino, 2017; Poirier et al., 2018; Tassetto et al., 2019; Zhu et al., 2022), it follows that its efficient intercellular spread (or that of derived signals, such as piRNAs or secondary siRNAs; Tassetto, Kunitomi, & Andino, 2017; Tassetto et al., 2019) might contribute to an efficient immune response in the insect body. In this scope, extracellular vesicles (EVs) deserve particular attention. EVs are membranous nanostructures, secreted by all known pro-and eukaryotic cell types, that can carry various regulatory molecules such as proteins, RNAs, and DNAs (Gill, Catchpole, & Forterre, 2019; van Niel et al., 2022; Remans et al., 2025). While insect EVs have been shown to contain potentially immunomodulatory RNA species (Kim et al., 2015; Lefebvre et al., 2016; Tassetto, Kunitomi, & Andino, 2017; Van den Brande et al., 2018; Tsai et al., 2019; Mingels et al., 2020), whether they can encompass vDNA is still to be determined.

Here we investigated vDNA formation in five insect species, either *in vivo* or *in vitro*. We used the non-retroviral cricket paralysis virus (CrPV), which has a +ssRNA genome, to investigate vDNA formation upon infection in various species belonging to the orders of Lepidoptera, Diptera, and Hemiptera. In addition, we assessed the presence of vDNA from persistently present non-retroviral RNA viruses, namely B. mori latent virus (BmLV; previously known as B. mori macula-like virus, MLV), Flock House virus (FHV), and Euscelidius variegatus virus 1 (EVV-1), all with a +ssRNA genome; and Halyomorpha halys toti-like virus 2 and H. halys partiti-like virus 1, both with a dsRNA genome. We also investigated the presence of CrPV-derived vDNA in EV preparations from infected S2 (*D. melanogaster*-derived) cells. Together, our results point towards a common mechanism of vDNA formation in insects, one that can be triggered upon a single infection with various RNA viruses and which allows secretion in association with EVs.

## Material and methods

### Cell culture

The lepidopteran High Five and BmN4 cells, derived from *Trichoplusia ni* and *B. mori*, respectively, were maintained in a complete medium consisting of IPL-41 Insect Medium (Sigma-Aldrich), 10% heat-inactivated fetal bovine serum (FBS) (Sigma-Aldrich), 0.25 μg/mL of amphotericin B (Sigma-Aldrich), 100 U/mL penicillin, and 100 μg/mL streptomycin (Thermo Fisher Scientific). The cells were maintained at 27.5 °C and sub-cultured weekly (1:10 and 1:5 respectively). The dipteran Schneider 2 (S2) cell line, derived from *D. melanogaster*, was maintained in a complete medium consisting of Shields and Sang M3 Insect medium (United States Biological Life Sciences) supplemented with 1 g/L yeast extract (Sigma-Aldrich) and 2.5 g/L bactopeptone (BD Biosciences), 10% heat-inactivated FBS (Sigma-Aldrich), 0.25 µg/mL Amphotericin B (Sigma-Aldrich), 100 U/ml penicillin, and 100 μg/ml streptomycin (Thermo Fisher Scientific). The cells were sub-cultured weekly (1:10) and maintained at 25 °C. FBS was heat-inactivated for 30 minutes at 56 °C prior to use.

### Insects

An *Euscelidius variegatus* (Kirschbaum) (Hemiptera, Cicadellidae) laboratory population (EvaTO) was originally collected in Piedmont (Italy). Insects were reared on oat, *Avena sativa* (L.), inside plastic and nylon cages in growth chambers at 20–25 °C with an L16:D8 photoperiod.

*Halyomorpha halys* (Stål) (Hemiptera, Pentatomidae) specimens were collected from Italian fields in 2021 within the framework of a project aiming at the virome characterization of natural populations (Papa et al., 2023). Specimens were stored in RNA*later*^TM^ (Thermo Fisher Scientific) upon collection and kept at −80°C until further processing.

### Production and quantification of CrPV

The CrPV suspension was produced in S2 cells as previously described (Garrey et al., 2010) and purified by ultracentrifugation on a sucrose cushion. The final viral pellet was resuspended in phosphate-buffered saline (PBS) or Tris buffer for the *in-vitro* and *in-vivo* experiments, respectively. For *in-vitro* experiments, the viral concentration was determined by transmission electron microscopy, with negative staining, by Sciensano (Ukkel, Belgium). For *in-vivo* experiments, a 1:1000 dilution of resuspended viral pellet was used.

### Infection with CrPV

*In-vitro* viral infections were performed as previously reported (Santos et al., 2018; Santos et al., 2019; Santos et al., 2022). Shortly, cells were pelleted in conical tubes for 10 minutes at 500x g and resuspended in basal medium containing CrPV suspension or PBS as a control (6 mL for every 2 x 10^6^ High Five or BmN4 cells, 6 mL for every 3 x 10^6^ S2 cells for detection of intracellular vDNA, and 6 mL for every 1.8 x 10^7^ S2 cells for EV isolation). 25 viral particles per cell were administered to High Five and BmN4 cells, and 15 viral particles per cell were administered to S2 cells. Afterwards, an incubation of 2 hours at room temperature on a shaker plate took place, followed by washing one time in basal medium and resuspension in complete medium (1 mL per 1 x 10^6^ High Five or BmN4 cells, 1 mL per 1.5 x 10^6^ S2 cells used for intracellular vDNA detection, and 1 mL per 7.2 x 10^5^ S2 cells for EV isolation). Basal medium refers to IPL-41 Insect Medium (Sigma-Aldrich) for High Five and BmN4 cells and to Shields and Sang M3 Insect medium (United States Biological Life Sciences) for S2 cells. Cells were then incubated as described in the section “Cell culture”. They were collected in 200µl of PBS and stored at −80°C until further processing.

CrPV infections of *E. variegatus* adults were achieved by intra-abdominal microinjection. Newly emerged adults were CO_2_-anesthetized and microinjected with 1 μl of CrPV virions resuspended in Tris buffer (10mM Tris buffer, pH 7.4) (1:1000 dilution of resuspended viral pellet). Mock-treated insects were microinjected with the same Tris buffer. Treated insects were checked daily for estimating the survival rate, and living insects were collected at 4 days post-injection (T4) and at 9 days post-injection (T9) for viral diagnosis by PCR analysis.

### EV isolation

The supernatants of CrPV-infected S2 cells were collected either immediately after infection (T0) or after 48 h (*i.e*., 2 days) of incubation (T2). The supernatants of mock-infected S2 cells were collected after 48 h of incubation (T2). All supernatants were first cleared of cells and apoptotic bodies through ten-minute centrifugations (4°C, 1000 g and 5000 g, respectively). The supernatants were then concentrated to 1 mL each using Amicon 10 kDa MWCO centrifugal filter units (Merck), and the EVs were isolated from the concentrated samples using size exclusion chromatography (SEC). SEC was performed in glass Econo-columns (#7374152; Bio-Rad) filled to a height of 14 cm - approximately 25 mL - with CL-6B Sepharose beads (Cytiva). The 1-mL samples were placed on top of the bead bed and allowed to elute in PBS through the column under the influence of gravity. 1-mL fractions were collected manually from the bottom of the column. Based on previous research (Van den Brande et al., 2025), fractions 8 and 9 were expected to contain EVs, and these fractions were combined prior to concentration to ca. 75 µL using Amicon 10 kDa MWCO centrifugal filter units (Merck).

### DNA extraction

DNA extraction from cell cultures was performed with the DNeasy Blood & Tissue Kit (Qiagen), following the manufacturer’s instructions (Purification of Total DNA from Animal Blood or Cells (Spin-Column Protocol), including RNase digestion). DNA from single adult insects was extracted using Direct-zol RNA Mini Prep kit (Zymo Research), following the manufacturer’s protocol, excluding the DNAse treatment and thereby allowing extraction of both DNA and RNA. The quality and concentration of the extracted DNA were assessed using a Nanodrop spectrophotometer (Thermo Fisher Scientific).

DNA was extracted from the EVs according to the protocol of Spada, Rudqvist, & Wennerberg (2020). In brief, the EV preparations were digested overnight at 56°C in a buffer containing 0.5% SDS, 50 mM Tris-HCl, 0.1 M EDTA (all from Merck), and 0.1 mg/mL proteinase K (Roche). Following the overnight digestion, DNA was extracted with phenol/chloroform (Merck) and precipitated with isopropanol. Quantification of DNA concentration and purity occurred with the Qubit High Sensitivity dsDNA Assay Kit (Thermo Fisher Scientific) on a Qubit® 2.0 Fluorometer (Thermo Fisher Scientific). Each sample was measured three times.

### DNase digestion

A DNase digestion of the purified DNA was performed for specific experimental set-ups. To this end, we used Amplification Grade DNase I (Deoxyribonuclease I) (Sigma-Aldrich) according to the manufacturer’s instructions.

### RNA extraction and cDNA synthesis

For RNA extraction of cultured cells, the miRNeasy Mini Kit (Qiagen) was used as described in the corresponding protocol. For extracting RNA to be used as a control for the EV-derived preparations, the RNeasy Mini Kit (Qiagen) was used, followed as necessary by cDNA synthesis (400 ng RNA per reaction; Takara PrimeScript Kit, Takara Bio). For all RNA extractions of cells, DNase treatment (RNase-free DNase set, Qiagen) was included in the workflow.

RNA from single adult insects was extracted using the Direct-zol RNA Mini Prep kit (Zymo Research), following the manufacturer’s protocol. Quality and concentration of the extracted RNA were assessed using a Nanodrop spectrophotometer (Thermo Fisher Scientific). Aliquots of RNA samples extracted from whole insects were DNase-treated (TURBO DNA-*free*™ Kit; Thermo Fisher Scientific). Then, cDNA was synthesized from total RNA samples (400 ng) using a High-Capacity cDNA Reverse Transcription Kit (Thermo Fisher Scientific) according to the manufacturer’s instructions.

### Polymerase chain reaction and gel electrophoresis

PCR was performed with sequence-specific primers (Table 1). Viral nucleotide sequences were retrieved from NCBI, the accession numbers being AF218039.1, NC_015524.1, EF690537.1, EF690538.1, OR074993.1, OR074979.1, and NC_032087.1. For the experiments involving cultured cells and derived EVs, REDTaq mix (Sigma-Aldrich) was used as a source of DNA Taq polymerase, dNTPs and PCR buffer. The following PCR program was used: 1x [10 minutes at 96oC]; 40x (35-50x for the EV-derived DNA) [1 minute at 96oC, 1 minute at the annealing temperature (Table 1), 30-45 seconds at 72oC]; 1x [5 minutes at 72oC]. As controls, we employed RNA derived from infected cells (a control for reverse-transcriptase activity of Taq polymerase; Jones & Foulkes, 1989), distilled water (a negative control), and cDNA derived from infected cells (a positive control).

**Table 1:**
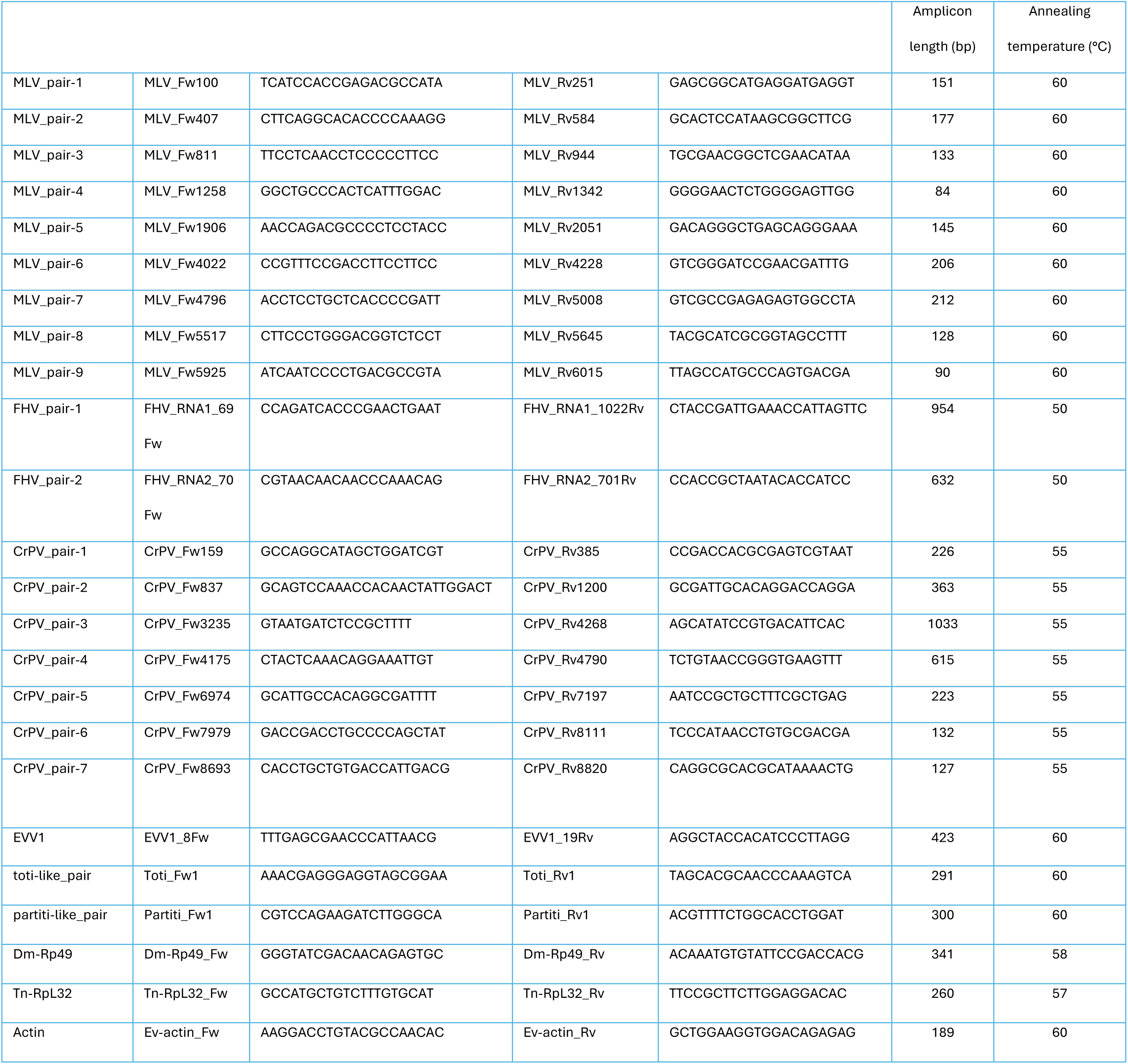
Sequences of the used primer pairs (5’-3’), respective amplicon lengths, and annealing temperatures. Fw – forward; Rv – reverse; bp – base pairs.

Genomic DNA from insect samples was amplified via PCR using AMPIGENE® Taq DNA Polymerase (Enzo Biochem Inc.). The following PCR program was used: 1x [1 minute at 94°C]; 35x [10 seconds at 96°C, 30 seconds at the primer annealing temperature (Table 1), and 30 seconds at 72°C]; 1x [5 minutes at 72°C]. As controls, we employed RNA subjected to DNase treatment derived from infected insects, distilled water as a non-template control (NTC), and cDNA derived from infected insects as a positive control.

The amplification products were subsequently analyzed by agarose gel electrophoresis with GelRed Nucleic Acid Gel Stain (Biotium) or SYBR Safe DNA Gel Stain (Thermo Fisher Scientific) and visualized under UV light. For estimation of amplicon lengths, DNA ladders were run along the samples, as indicated in the figure captions.

### Amplicon sequence confirmation

The identity of the obtained bands was always confirmed by Sanger sequencing. After electrophoresis, individual bands were cut out of the gel, and the DNA was extracted with the GenElute Gel Extraction Kit (Sigma-Aldrich) or the Wizard® SV Gel and PCR Clean-Up System (Promega). This DNA served as template for a second PCR round as described above, followed by a cleanup with the GenElute PCR Clean-Up Kit (Sigma-Aldrich). This product was then either directly Sanger-sequenced (LGC Genomics) or cloned into a pGem-T Easy Vector (Promega) and then Sanger-sequenced (BMR Genomics).

## Results

### CrPV vDNA is formed in dipteran cells upon infection

We started by investigating if vDNA is detected in *D. melanogaster* S2 cells after CrPV infection. An initial test was performed in which cells were infected or mock-treated followed by DNA purification 24 hours later. Then a PCR analysis was performed by using seven different CrPV primer pairs, four of which yielded a positive result (**Figure S1, panel A**). With this result, we selected one primer pair to further test two control conditions. Specifically, to exclude that potential residual RNA could serve as a PCR template, RNA purified from the same conditions was tested. Furthermore, to confirm that DNA was in fact the PCR template, we also tested DNA digested with DNase. A band was observed only when intact DNA of infected cells was used as PCR template (**Figure S1, panel B**). Next, we performed the infection (or mock infection with PBS) and purified DNA at time zero (T0, immediately after infection) and at three days post-treatment (T3). Afterwards, we performed PCR with CrPV-specific primers (CrPV_pair-5) and analyzed the products by gel electrophoresis (**Figure 1**). When DNA purified three days after infection was used, a specific band was observed. This was not the case when DNA purified immediately after infection was used, or when the cells were mock-treated, showing that CrPV vDNA was formed upon viral infection and present at least until three days post-infection (**Figure 1**). Bands were observed for these samples when primers specific for a reference gene (*Dm-Rp49*) were used, confirming gDNA quality in every sample (**Figure S1, panel C).**

**Figure 1:**
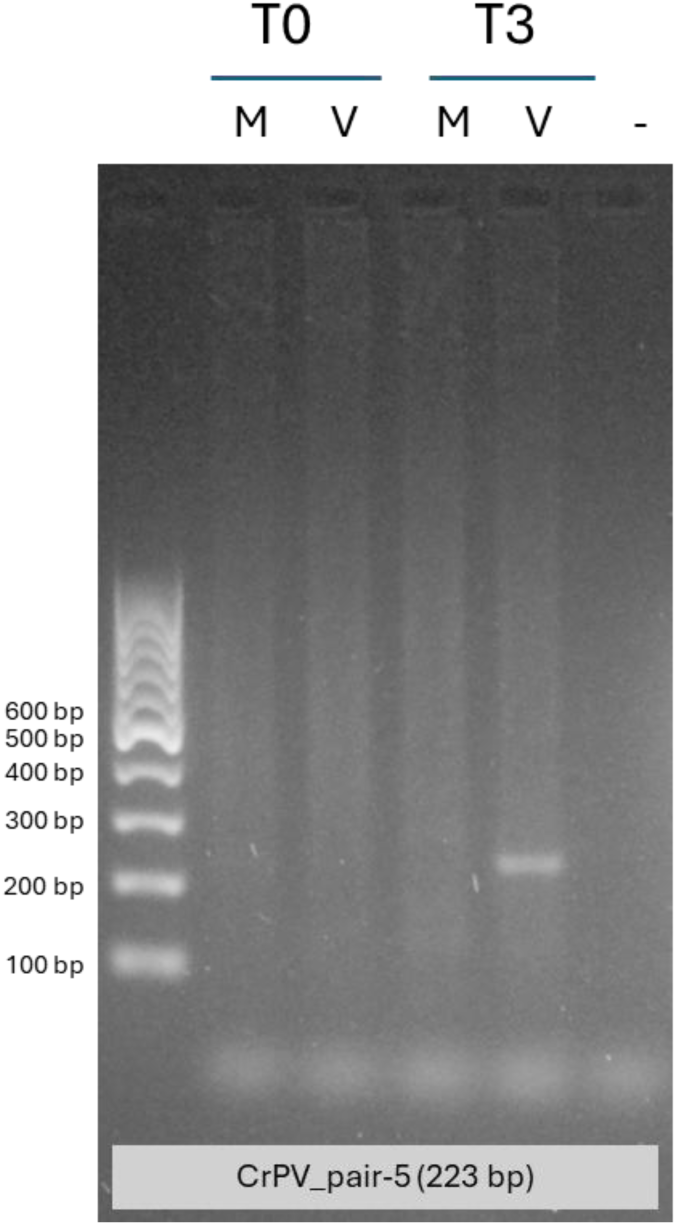
Presence of intracellular vDNA from CrPV in S2 cells upon infection. Amplicons produced with CrPV-specific primers (CrPV_pair-5) from cellular DNA extracts collected immediately (T0) or three days (T3) after viral infection (V) or mock-infection (M). NTC = non-template control. The Thermo Fisher Scientific GeneRuler 100 bp DNA ladder was used as a reference.

### CrPV vDNA is formed in lepidopteran cells upon infection

We then investigated if vDNA is formed in the *T. ni* High Five cell line upon infection with CrPV. To this end, we performed the infection (or PBS mock-infection) and purified DNA at time zero (T0, immediately after infection) and at three days post-treatment (T3). Afterwards, we performed PCR with CrPV-specific primers (CrPV_pair-5), and the products were analyzed by gel electrophoresis (**Figure 2**). When DNA purified three days after infection was used, a specific band was observed. This was not the case when DNA purified immediately after infection was used, or when the cells were mock-treated, showing that CrPV vDNA was formed upon viral infection and present at least until three days post-infection (**Figure 2**). Bands were observed for these samples when primers specific for a reference gene (*Tn-RpL32*) were used, confirming gDNA quality (**Figure S2**). To exclude that potential residual RNA could serve as PCR template, RNA purified from the same conditions was tested. Furthermore, to confirm that DNA was in fact the PCR template, we also tested DNA digested with DNase. No bands were observed in these conditions (**Figure S2**).

**Figure 2:**
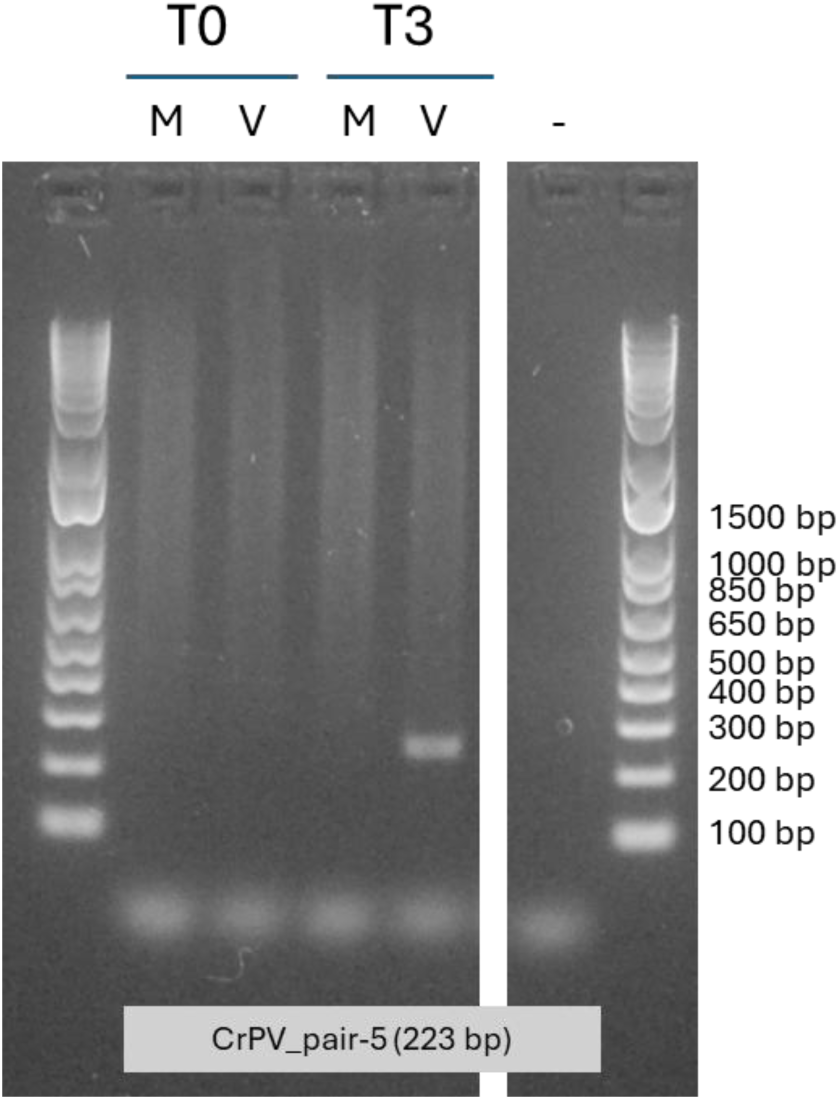
Presence of intracellular vDNA from CrPV in High Five cells upon infection. Amplicons produced with CrPV-specific primers (CrPV_pair-5) from cellular DNA extracts collected immediately (T0) or three days (T3) after viral infection (V) or mock infection (M). -: non-template control. The Thermo Fisher Scientific 1 Kb Plus DNA ladder was used as a reference.

Afterwards, we investigated if vDNA can also be detected in *B. mori* BmN4 cells after CrPV infection. To this end, cells were infected or mock–treated with PBS followed by DNA purification three days later. An initial test was performed by using three different CrPV primer pairs, two of which yielded a positive result (**Figure S3, left panel**). We then selected one primer pair (CrPV_pair-5) and showed that a specific band was observed when DNA purified three days after infection was used as PCR template, while this was not the case when DNA purified immediately after infection was used, or when the cells were mock-treated (**Figure 3**). We also tested two control conditions, namely RNA and DNA digested with DNase, as explained above. A band was observed only when undigested DNA of infected cells was used as PCR template, confirming that CrPV vDNA was formed and present at least until three days post-infection in these cells (**Figure S3, right panel**).

**Figure 3:**
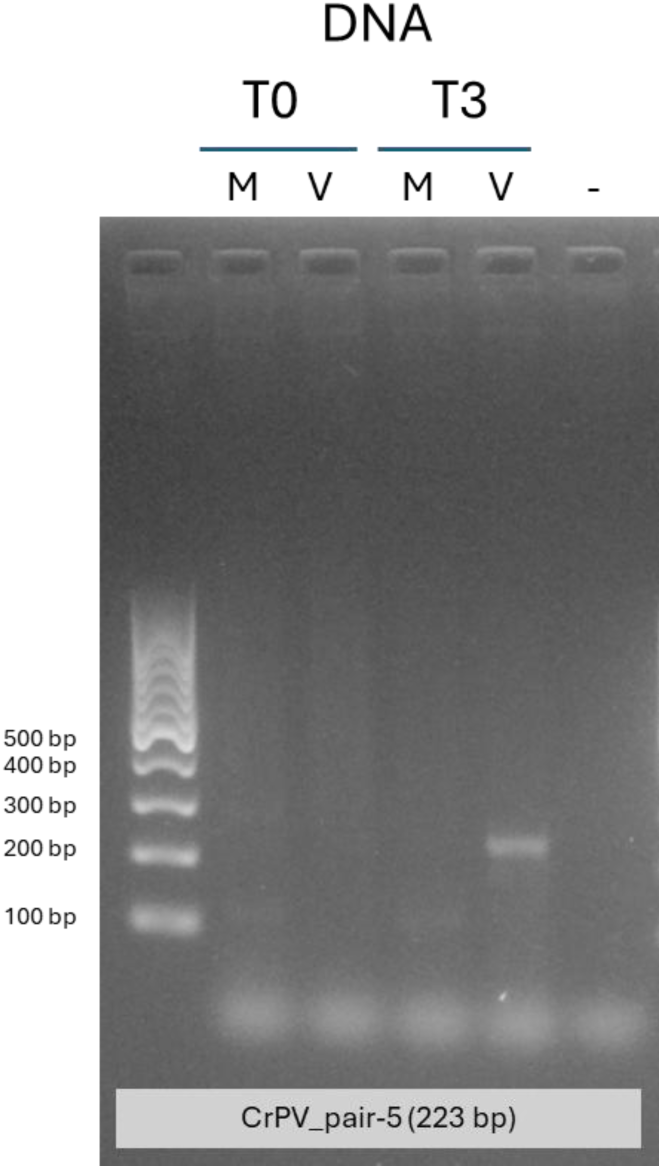
Presence of intracellular vDNA from CrPV in BmN4 cells upon infection. Amplicons produced with CrPV-specific primers (CrPV_pair-5) from cellular DNA extracts collected immediately (T0) or three days (T3) after viral infection (V) or mock infection (M). -: non-template control. The Thermo Fisher Scientific GeneRuler 100 bp DNA ladder was used as a reference.

### CrPV vDNA is formed in *E. variegatus* insects upon infection

Next, we investigated if vDNA was also formed upon CrPV infection in the leafhopper, *E. variegatus.* To this end, *E. variegatus* adults were microinjected with CrPV virions (or a mock infection) followed by extraction of total nucleic acids at 4 and 9 days post-injection (4 dpi and 9 dpi, respectively). An initial PCR test was performed by using four different CrPV primer pairs (CrPV_pair-1, −5, −6, and −7), one of which yielded a positive result (CrPV_pair-1). Specifically, with this primer pair, a CrPV-specific band was observed when DNA collected 4 and 9 days post-injection was amplified (**Figure 4**). CrPV-infected samples resulted in the expected 226-bp amplicon, while the mock treatment produced two off-target amplicons of ∼400bp and ∼900bp, respectively (**Figure 4**). By using DNA digested with DNase as PCR template, we also confirmed that DNA was, in fact, the template for the previously observed bands (**Figure S4**).

**Figure 4:**
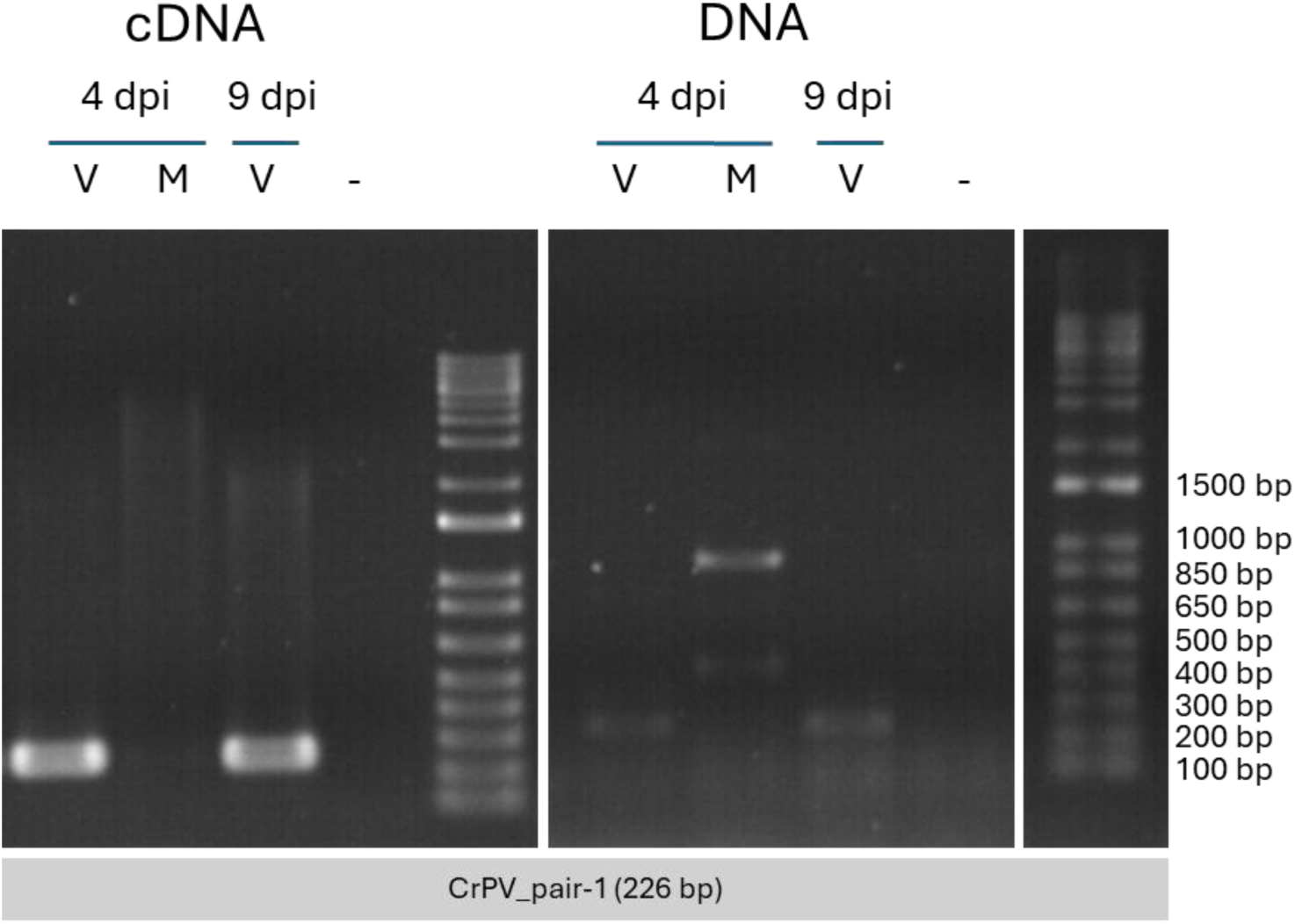
Presence of vDNA from CrPV in *E. variegatus* insects upon infection. Amplicons produced with CrPV-specific primers (CrPV_pair-1) from cDNA (positive control) or DNA collected four (4 dpi) or nine (9 dpi) days after viral infection (V) or mock infection (M). -: non-template control. The Thermo Fisher Scientific 1 Kb Plus DNA ladder was used as a reference.

### CrPV vDNA is associated with dipteran cell EVs

Having detected CrPV-derived vDNA in both cell lines and whole insects, we decided to investigate whether EVs from infected cells (the S2 cell line in this case) carry these vDNA sequences. Each primer pair was tested on the three EV-derived DNA samples described above in the Methods section (mock 48 hpi, CrPV 0 hpi, and CrPV 48 hpi).

For primer pairs 1, 5, 6, and 7, amplicons of the expected length appeared only in the positive control and in the virus-infected samples collected 48 hpi (T2) (**Figure 5**). Primer pair 4 resulted in nonspecific amplification in both the virus-infected T2 and the mock-infected T2 conditions; Sanger sequencing confirmed that this sequence originated from *D. melanogaster*. Amplicons of all primer pairs were absent from the samples collected immediately after viral infection (T0). Likewise, no amplification was detected in the RNA-only control reactions.

**Figure 5:**
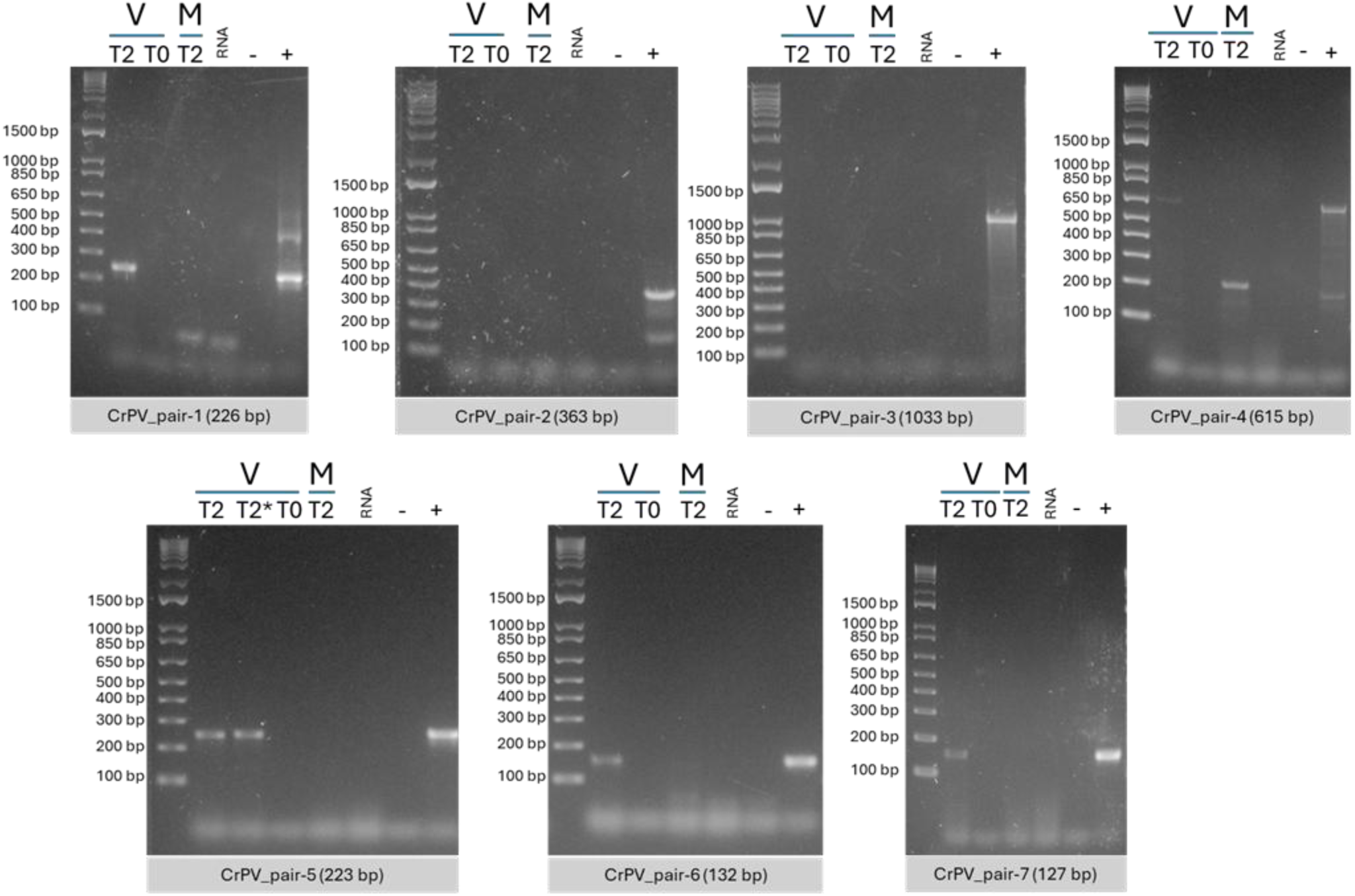
Presence of vDNA from CrPV in S2 EVs. Seven different primer pairs targeting the CrPV genome were used. Amplicons were produced with EV-extracted DNA from virus-infected (V) or mock-infected (M) cells immediately (T0) or 48 h/two days (T2) after treatment. RNA: RNA-only control; -: non-template control; +: positive control (cDNA). For CrPV_pair-5, one reaction (*) was performed with 7.5 ng DNA; the other reactions for this primer pair were set up with 15 ng DNA each. The Thermo Fisher Scientific 1 Kb Plus DNA ladder was used as a reference.

### vDNA of persistent viral infections is present in lepidopteran cells

The lepidopteran cell lines used in this study are persistently infected with BmLV (High Five and BmN4 cells) and/or with FHV (High Five cells) (*e.g.* Katsuma et al., 2005; Santos et al., 2022; Swevers et al., 2016; Verdonckt et al., 2023). To test if vDNA for these non-reverse transcribing (non-RT) RNA viruses is present under these circumstances, we first purified DNA from these cells and performed PCR with virus-specific primers (**Figure S5**). For the BmN4 cells, this PCR resulted in a specific band for eight out of nine tested BmLV-specific primers (**Figure S5 A**). For the High Five cells, the PCR resulted in a specific band for eight out of nine BmLV-specific primer pairs (**Figure S5 B**) and a specific band for two FHV-specific primer pairs (**Figure S5 C**). Based on this result, we selected one primer pair for each virus to further test two control conditions, namely RNA and DNA digested with DNase, as explained above. A band was observed only when undigested DNA was used as a PCR template, confirming that BmLV vDNA is present in BmN4 cells, and that vDNA from both BmLV and FHV is present in High Five cells (**Figure 6**).

**Figure 6:**
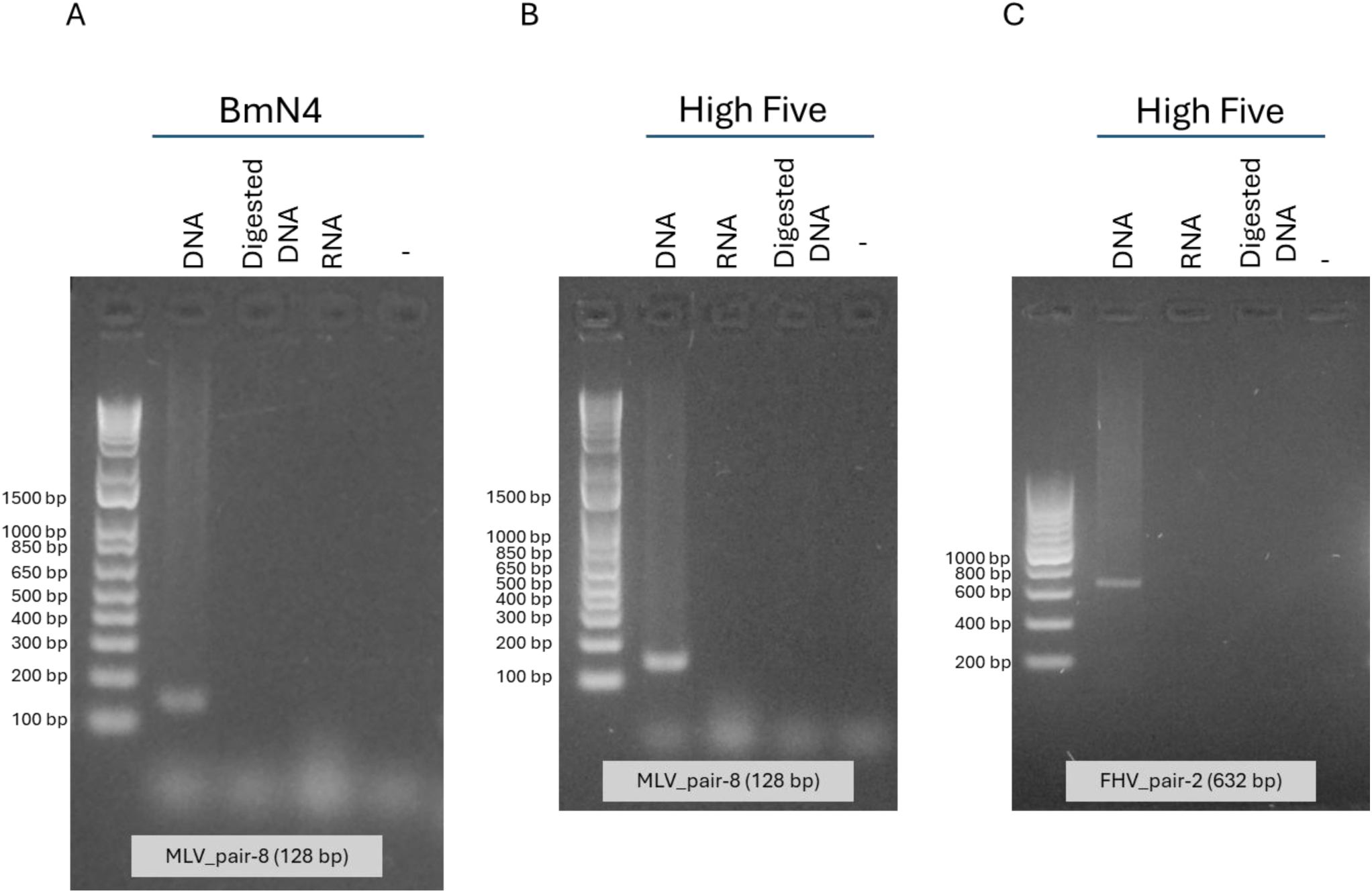
vDNA from persistently present non-RT RNA viruses in lepidopteran cell lines. **A:** Amplicons produced with BmLV-specific primers (MLV_pair-8) from DNA, digested DNA, and RNA from *B. mori* BmN4 cells persistently infected with BmLV. **B:** Amplicons produced with BmLV-specific primers (MLV_pair-8) from DNA, RNA, and digested DNA from *T. ni* High Five cells persistently infected with BmLV and FHV. **C:** Amplicons produced with FHV-specific primers (FHV_pair-2) from *T. ni* High Five cells persistently infected with BmLV and FHV. -: non-template control. The Thermo Fisher Scientific 1 Kb Plus DNA Ladder was used as a reference for panels A and B, while the Thermo Fisher Scientific O’Range Ruler 200 bp DNA Ladder was used as a reference for panel C.

### vDNA of persistent viral infections is present in *E. variegatus* insects

The EvaTO laboratory population was previously found to be infected by EVV1, a well-characterized iflavirus causing a persistent and vertically transmitted covert infection with a 100% prevalence in this population (Abbà et al., 2017; Ottati et al., 2020). Nine *E. variegatus* insects persistently infected with EVV1 were used to test if vDNA for this non-RT RNA virus would be present under these circumstances. Total nucleic acids were extracted from nine EVV1- infected insects, which were then tested through PCR with virus-specific primers (**Table 1**). We observed amplification of EVV1 vDNA from DNA of six individuals and none from DNAse-digested RNA or cDNA (**Figure 7**). Sequencing of the obtained amplicons confirmed the presence of EVV1 vDNA in EVV1-infected insects. Bands were observed for all samples when primers specific for a reference gene (*actin*) were used, confirming gDNA quality (**Figure S6**).

**Figure 7:**
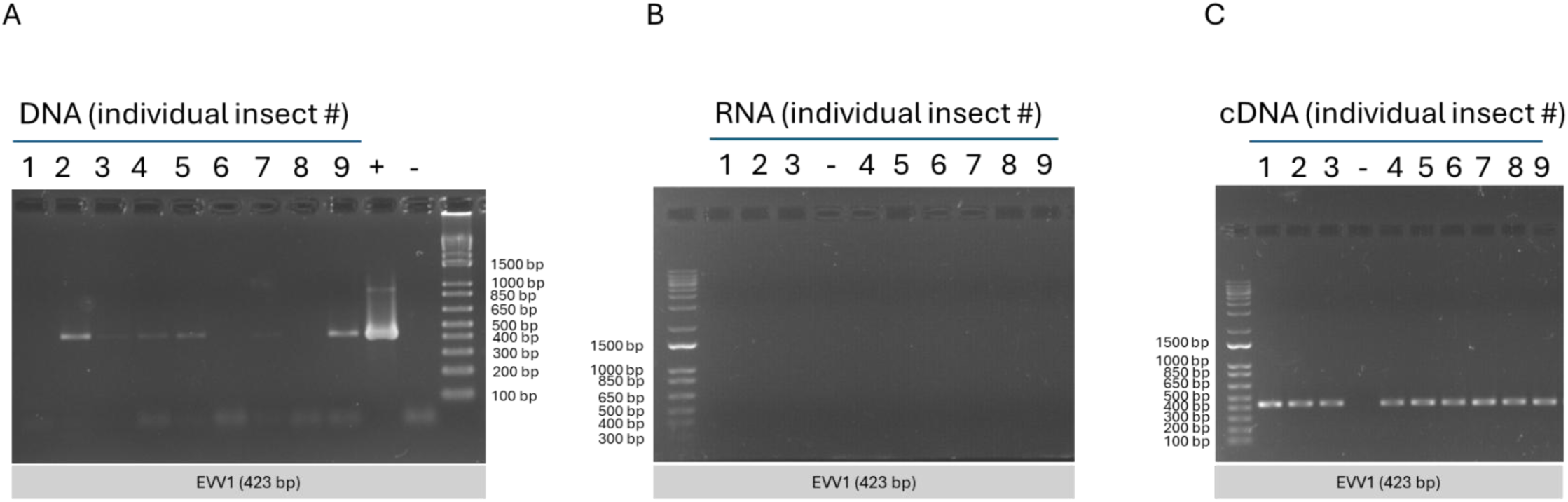
vDNA of persistently present EVV1 in *E. variegatus* individuals. Amplicons produced by PCR with EVV1-specific primers from DNA of persistently infected individuals, a positive control (+ control) represented by the cDNA of one EVV1-infected insect, and an NTC (non-template control). **A:** EVV1-specific primers were used to amplify DNA. **B:** EVV1-specific primers were used to amplify RNA aliquots (treated with DNase) and (**C**) cDNA. The Thermo Fisher Scientific 1 Kb Plus DNA ladder was used as a reference.

### vDNA of newly identified viruses is present in field-sampled *H. halys* insects

Field-collected *H. halys* samples infected with H. halys toti-like virus 2 or with H. halys partiti-like virus 1, both non-RT dsRNA viruses, were used to test vDNA presence. We conducted an initial test by purifying total nucleic acids from adult insects and performing PCR with virus-specific primers (Table 1) on the insect cDNA and DNA to confirm the infection and to detect the vDNA, respectively (**Figure 8**). For RNA-only controls, we digested the total nucleic acids extracted from *H. halys* individuals with DNase and used the remaining RNA as PCR templates. For both viruses, amplification of vDNA was observed from both insect DNA and cDNA, but not from RNA digested with DNAse. Sequencing of the amplicons confirmed the presence of vDNA of H. halys toti-like virus 2 and H. halys partiti-like virus 1 in the infected insects.

**Figure 8:**
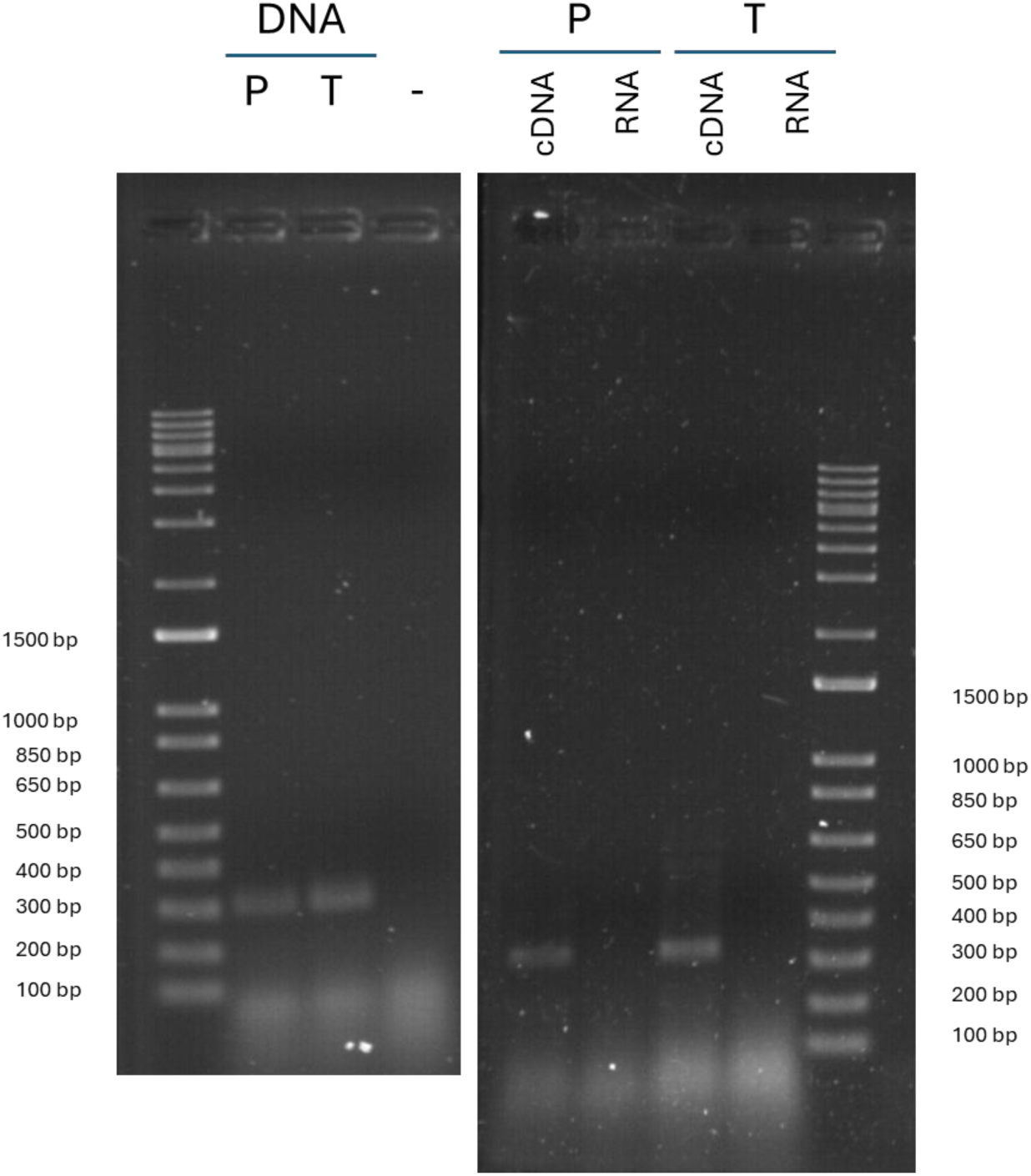
Detection of viral DNA (vDNA) of persistently present H. halys partiti-like virus 1 (**P**) and H. halys toti-like virus 2 (**T**) in *H. halys* individuals. Left: DNA extracted from persistently infected individuals was used as a template. Right: cDNA and RNA extracted from persistently infected individuals were used as templates. -: non-template control. The Thermo Fisher Scientific 1 Kb Plus DNA ladder was used as a reference.

## Discussion

In this study, we found vDNA sequences mapping to the viral genomes of several non-reverse transcribing RNA viruses in various insects and insect-derived cell lines. Specifically, we found that infection with CrPV results in *de novo* formation of vDNA in S2 cells, and that this vDNA is secreted in association with EVs. We also found that vDNA is produced in response to infection in lepidopteran cell lines, as well as in hemipteran insects.

Previous studies detecting vDNA formation in Diptera did so in whole flies and mosquitoes, the embryo-derived Aag2 and C6/36 mosquito cell lines, and the haemocyte-like S2 cells (Goic et al., 2013; Goic et al., 2016; Nag, Brecher, & Kramer, 2016; Poirier et al., 2018; Tassetto, Kunitomi, & Andino, 2017; Mondotte et al., 2018; Tassetto et al., 2019; Mondotte et al., 2020; Uddin et al., 2024). In Lepidoptera, vDNA has been reported from the BmN cell line (Zhu et al., 2022). VDNA formation has, to our knowledge, never been reported in Hemiptera or Lepidoptera outside *B. mori*. Therefore, our discoveries in *T. ni* cells, *E. variegatus*, and *H. halys* broaden the range of species in which viral infection-induced vDNA formation has been described. The species we examined belong to evolutionarily distinct lineages, suggesting that these observations reflect a widespread mechanism in insects. Our use of cell lines of different tissue origins also suggests that the formation of vDNA from non-retroviral RNA viruses may be a general feature of (at least some) cells within the insect body. Both BmN4 and High Five cells are ovary-derived, yet BmN4 cells are described to be derived from the germline (Honda et al., 2017), while High Five cells are similar to S2 cells in displaying hemocyte-like characteristics (Beck & Strand, 2005). We also showed that a wide range of viruses can induce vDNA formation. Although the S2 cell line is known to produce vDNA when exposed to Flock House, Drosophila X, and Drosophila C viruses (Goic et al., 2013; Poirier et al., 2018), its response to CrPV has not been investigated before. Likewise, the BmN cell line (from which our BmN4 cells were derived) was previously shown to produce vDNA in response to B. mori cytoplasmic polyhedrosis virus (Zhu et al., 2022), a dsRNA virus; we showed that these *B. mori* cells also produce vDNA when infected with the dicistrovirus CrPV and the tymovirus BmLV, both RNA viruses.

Prior studies in mosquitoes suggested that vDNA may integrate itself into the genome, thereby forming EVEs over time (Rodriguez-Andres et al., 2024; Tassetto et al., 2019; Uddin et al., 2024). Those observations suggest that viral DNA sequences in insect genomes may represent genomic records of past infections. Our study detected virus-specific DNA produced *de novo* upon (recent) viral infection, instead of constituting an established element of the insects’ or cells’ genomes. For CrPV-infected cells and insects, vDNA was not observed in mock-infected or T0 conditions (**Figures 1-4**), indicating that it was formed after infection. Likewise, EVs from S2 cells were CrPV vDNA-positive 48 hpi, but not immediately after infection (T0) or 48 hours after mock infection (**Figure 5**). The EV-associated DNA, therefore, is not a general feature of EVs that are produced by S2 cells or that are already present in the serum that was added to the cell medium. Notably, some of the viruses we examined have established persistent infections in their hosts (**Figures 6-8**), raising the possibility that the vDNA we detected might have been derived from EVEs integrated into the genome. However, sequencing showed that our amplicons closely matched existing viral sequences, while EVEs typically show substantial divergence with existing viruses (Suzuki et al., 2020; Uddin et al., 2024). Our results for the hemipteran insects further suggest that the vDNA was not a common genomic feature. For *E. variegatus*, not all individuals that we sampled tested positive for vDNA of their respective viruses (**Figure 7**). Had amplification of EVEs (*i.e.*, established genomic sequences) occurred, all of the individuals would have tested positive for the DNA sequences in question. For *H. halys*, RNA sequencing showed that not all collected individuals tested positive for the viruses under consideration (Papa et al., 2023), and only those insects that did test positive exhibited vDNA sequences (**Figure 8**).

In flies, mosquitoes, and silkworms, vDNA formation has been shown to enable resistance to subsequent viral infections (Tassetto, Kunitomi, & Andino, 2017; Mondotte et al., 2018; Poirier et al., 2018; Tassetto et al., 2019; Mondotte et al., 2020; Zhu et al., 2022; Rodriguez-Andres et al., 2024). Despite these links with antiviral immunity, the function of vDNA in the class of insects is not entirely clear. We observed that vDNA formation occurred quite rapidly after infection, as early as 24 hpi in the case of S2 cells. In fact, previous studies found that vDNA formation can occur as early as 6 hpi in the case of *Aedes albopictus* cell lines infected with Chikungunya virus (Goic et al., 2016). We also noted that CrPV vDNA remained detectable up to nine days post-infection in the case of *E. variegatus* (**Figure 4**). Again, this observation is in line with previous studies, some of which demonstrated that vDNA persists after clearance of viral infection (Mondotte et al., 2018) and can even be transmitted to and enable antiviral resistance in the insect’s progeny (Mondotte et al., 2020; Rodriguez-Andres et al., 2024). In *D. melanogaster*, evidence showed that vDNA can be transcribed into secondary siRNAs by the hemocytes, apparently through the action of endogenous reverse transcriptases (Tassetto, Kunitomi, & Andino, 2017). Flies thus appear to utilize vDNA to amplify their RNAi responses in the absence of RNA-dependent RNA polymerases (RdRPs), which produce secondary siRNAs in plants and nematodes, but which have not been found in insects (Carthew & Sontheimer, 2009). vDNA also appears to enable a certain degree of immune memory in flies, possibly even mediating the transmission of antiviral resistance to progeny (Mondotte et al., 2020; Rodriguez-Andres et al., 2024). However, the role of vDNA in immune memory remains to be proven beyond doubt.

Given its potential link with immunity, efficient spread of vDNA could be beneficial for the insect. Transport via EVs could be one possible means of accomplishing this spread of vDNA. In insects and other arthropods, small (s)RNAs – components of the RNAi response – have been detected several times in association with EVs (Tassetto, Kunitomi, & Andino, 2017; Van den Brande et al., 2018; Tsai et al., 2019; Yang et al., 2019; Mingels et al., 2020; Nawaz et al., 2020). In this study, we showed that insect EVs also carry, or at least are associated with, vDNA. Although DNA has been detected multiple times in association with mammalian EVs (Dixson et al., 2023), its presence in or on insect EVs has not been investigated much. Kaur et al. (2025) mentioned the presence of DNA in *B. mori* EV preparations but did not further elucidate the identity of these sequences. Studies in mammals suggest that DNA can be carried both inside EVs and attached to the EV surface (Hallal et al., 2022). In our study, we did not discriminate between the two options, as lumen-carried as well as surface-bound molecules can be part of the functional cargo repertoire of EVs (Hallal et al., 2022; Dixson et al., 2023). EV preparations may also contain various nonvesicular contaminants, including DNA (Jeppesen et al., 2019; Liangsupree, Multia, & Riekkola, 2021). However, the SEC-based protocol we used is considered one of the most highly efficient methods of removing contaminants from EV preparations, including those derived from insect cell cultures (Liangsupree, Multia, & Riekkola, 2021; Van den Brande et al., 2025).

EVs have been known for some time to carry out immunomodulatory signaling functions in insects, such as through the transport of dsRNA-derived sRNAs (Mingels et al., 2020). Tassetto, Kunitomi, & Andino (2017) further demonstrated that EVs may carry secondary siRNAs, transcribed from vDNA, in *D. melanogaster*. Considering these previous findings, our discovery suggests that EVs may act as intercellular messengers that circulate in the haemolymph to deliver vDNA to recipient cells. These recipient cells then may take up the vDNA to transcribe it into secondary siRNAs, in addition to or instead of receiving EV-delivered siRNAs directly (Tassetto, Kunitomi, & Andino, 2017; Mingels et al., 2020).

Based on our findings, it is now clear that multiple evolutionarily divergent insect species are capable of producing vDNA upon infection with diverse RNA viruses that lack encoded reverse transcriptases. These results raise important questions about the precise mechanisms driving vDNA generation, the molecular factors involved, and the functional significance of this process, particularly in the context of insect antiviral immunity. While detecting infection-induced vDNA formation represents a crucial first step, functional studies are now needed to test whether vDNA contributes to immune defences in insect orders such as Lepidoptera and Hemiptera. Likewise, the relevance of EV-mediated vDNA transfer to antiviral immunity remains to be established. Future mechanistic and functional research will be essential to elucidate the roles of vDNA produced during non-retroviral RNA virus infections, not only in fruit flies or mosquitoes, but also more broadly across insects, the most species-rich class of animals.

## Acknowledgements

The authors would like to thank Ilaria Negri and Giulia Papa for providing *Halyomorpha halys* specimens and Emilyn Matsumura for providing the CrPV infectious clone. The authors also wish to thank Annick Francis for her technical contributions to the experiments presented in this work.

---

**Figure S1:** Presence of intracellular vDNA from CrPV upon infection in S2 cells. **Panel A:** amplicons produced with CrPV-specific primers from DNA or RNA extracts of virus-infected (V) or mock-infected (M) cells. **Panel B:** Amplicons produced with CrPV-specific primer pair 7 from DNA, RNA, or digested DNA from virus-infected (V) or mock-infected (M) cells. **Panel C:** Amplicons produced with *Dm*Rp49-specific primers from DNA of cells collected immediately (T0) or three days (T3) after viral infection (V) or mock-infection (M). -: non-template control. The Thermo Fisher Scientific 1 Kb Plus DNA ladder was used as a reference for panels A and B, while the Thermo Fisher Scientific GeneRuler 100 bp DNA ladder was used as a reference for panel C.

**Figure S2:** Presence of intracellular vDNA from CrPV in High Five cells upon infection. **Left:** Amplicons produced with CrPV-specific primers (CrPV_pair-5) from DNA, RNA, or digested DNA, collected immediately (T0) or three days (T3) after viral infection (V) or mock-infection (M) of cells. **Right:** Amplicons produced with *Tn*RpL32-specific primers from DNA of cells collected immediately (T0) or three days (T3) after viral infection (V) or mock infection (M). -: non-template control. The Thermo Fisher Scientific 1 Kb Plus DNA ladder was used as a reference.

**Figure S3:** Presence of intracellular vDNA from CrPV in BmN4 cells upon infection. **Left:** Amplicons produced with CrPV-specific primers from DNA of virus-infected (V) or mock-infected (M) cells. **Right:** Amplicons produced with CrPV-specific primers (CrPV_pair-1) from DNA, RNA, or digested DNA of virus-infected (V) or mock-infected (M) cells. -: non-template control. The Thermo Fisher Scientific GeneRuler 100 bp DNA ladder was used as a reference.

**Figure S4:** Controls for the CrPV vDNA detected in *E. variegatus* insects. Amplicons produced with CrPV-specific primers (CrPV_pair-1) from digested DNA, collected four (4 dpi) or nine (9 dpi) days after viral infection (V) or mock infection (M). The Thermo Fisher Scientific 1 Kb Plus DNA ladder was used as a reference.

**Figure S5:** vDNA from persistently present non-RT RNA viruses in lepidopteran cell lines. **A:** Amplicons produced by PCR with BmLV-specific primers from DNA extracted from *B. mori* BmN4 cells persistently infected with BmLV. Left to right, MLV_pair-1-9 were used, as indicated. **B:** Amplicons produced with BmLV-specific primers from DNA from *T. ni* High Five cells persistently infected with BmLV and FHV. Left to right, MLV_pair-1-9 were used, as indicated. **C:** Amplicons produced with FHV-specific primers from DNA from *T. ni* High Five cells persistently infected with BmLV and FHV. Left to right, FHV_pair-1-2 were used, as indicated. The Thermo Fisher Scientific 1 Kb Plus DNA Ladder was used as a reference for panels S5A and S5B, while the Thermo Fisher Scientific O’Range Ruler 200 bp DNA Ladder was used as a reference for panel S5C.

**Figure S6:** Amplicons produced by PCR with *E. variegatus* actin-specific primers from DNA of *E. variegatus* individuals persistently infected with EVV1. The Thermo Fisher Scientific 1 Kb Plus DNA ladder was used as a reference.

