## Supplementary figures for "Infection with non-retroviral RNA viruses produces virus-derived DNA across diverse insect species"

### Slide 1
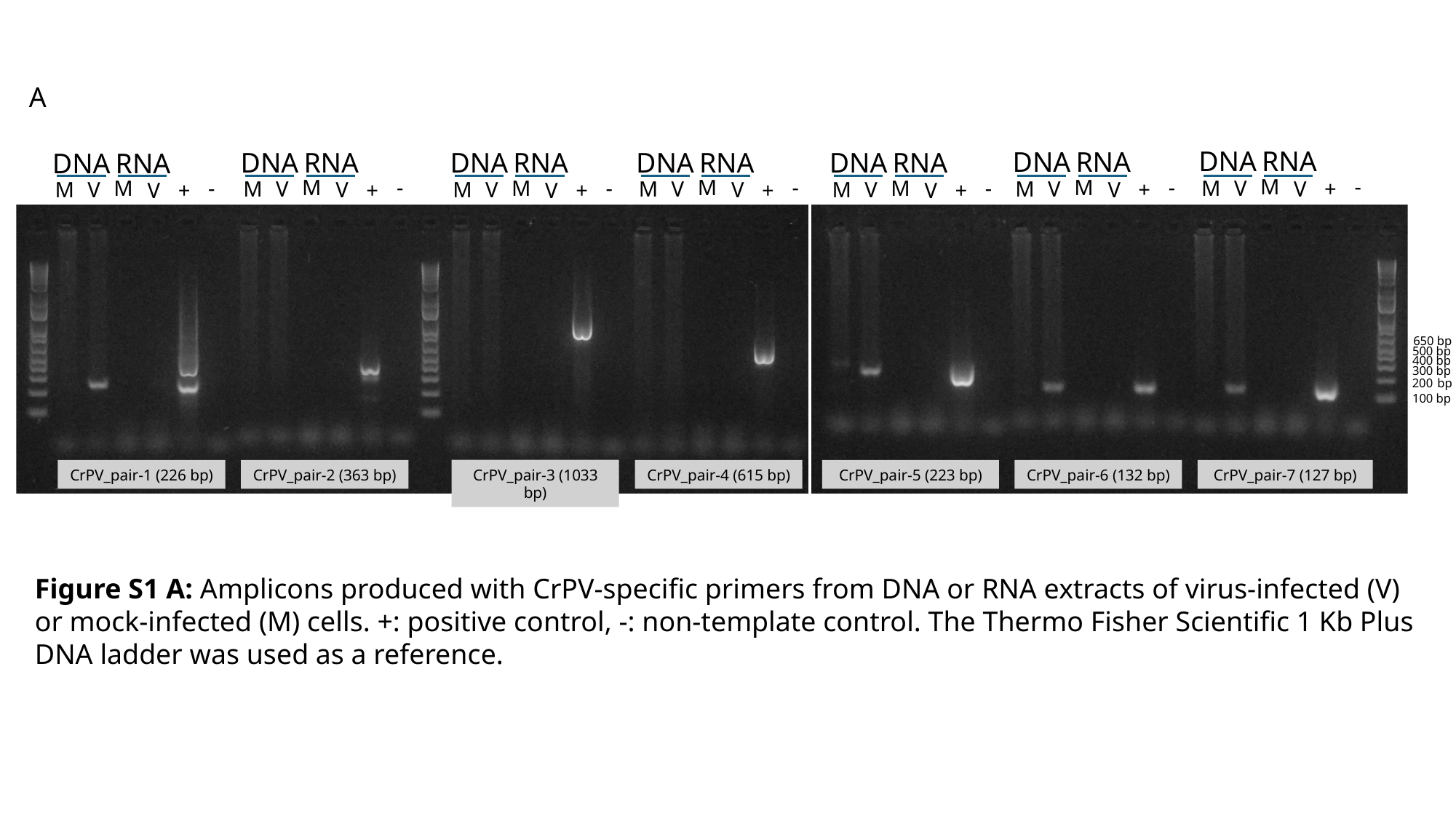

A
DNA
RNA
DNA
RNA
DNA
RNA
DNA
RNA
DNA
RNA
DNA
RNA
DNA
RNA
M
-
M
-
M
-
M
-
V
M
M
-
M
-
M
-
V
V
M
V
V
M
M
+
V
V
V
V
V
V
M
M
M
+
+
+
V
V
V
+
+
+
CrPV_pair-5 (223 bp)
CrPV_pair-7 (127 bp)
CrPV_pair-1 (226 bp)
CrPV_pair-2 (363 bp)
CrPV_pair-3 (1033 bp)
CrPV_pair-4 (615 bp)
CrPV_pair-6 (132 bp)
650 bp
500 bp
400 bp
300 bp
200 bp
100 bp
Figure S1 A: Amplicons produced with CrPV-specific primers from DNA or RNA extracts of virus-infected (V) or mock-infected (M) cells. +: positive control, -: non-template control. The Thermo Fisher Scientific 1 Kb Plus DNA ladder was used as a reference.

### Slide 2
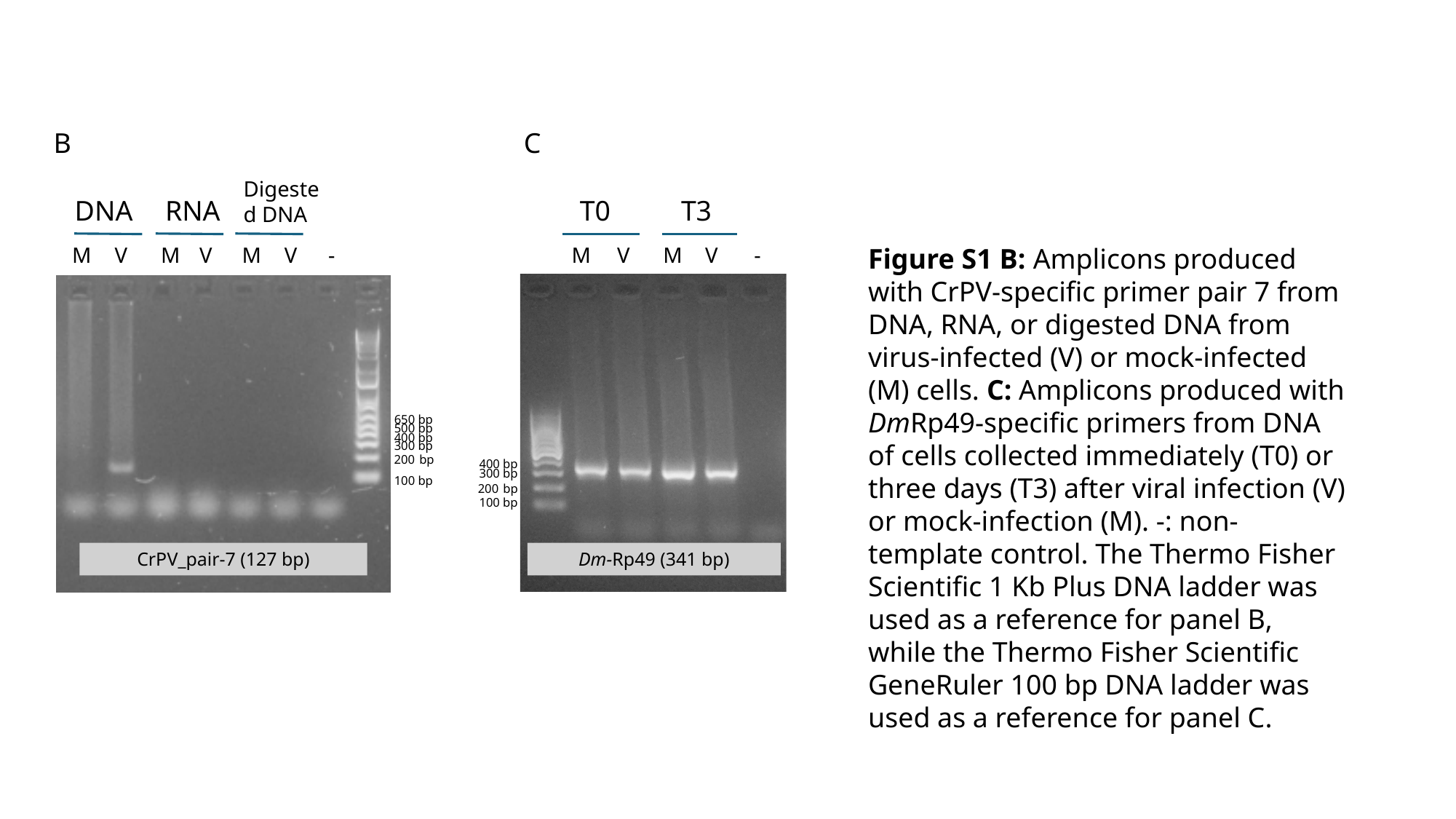

B
C
Digested DNA
DNA
RNA
T0
T3
M
V
M
V
M
V
-
M
V
M
V
-
Figure S1 B: Amplicons produced with CrPV-specific primer pair 7 from DNA, RNA, or digested DNA from virus-infected (V) or mock-infected (M) cells. C: Amplicons produced with DmRp49-specific primers from DNA of cells collected immediately (T0) or three days (T3) after viral infection (V) or mock-infection (M). -: non-template control. The Thermo Fisher Scientific 1 Kb Plus DNA ladder was used as a reference for panel B, while the Thermo Fisher Scientific GeneRuler 100 bp DNA ladder was used as a reference for panel C.
Dm-Rp49 (341 bp)
650 bp
500 bp
400 bp
300 bp
200 bp
400 bp
300 bp
100 bp
200 bp
100 bp
CrPV_pair-7 (127 bp)

### Slide 3
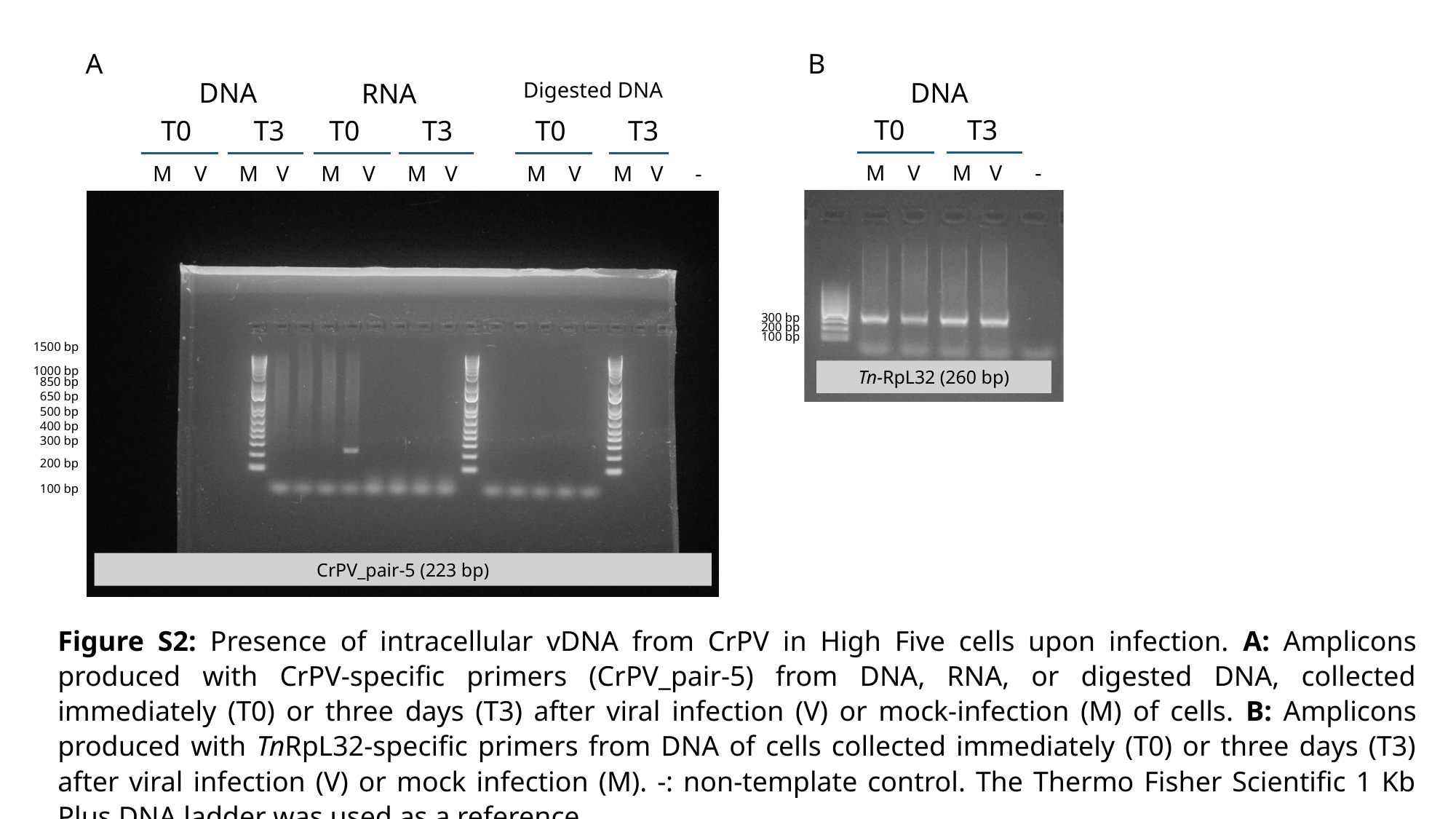

A
B
DNA
DNA
RNA
Digested DNA
T0
T3
T0
T3
T0
T3
T0
T3
M
V
M
V
-
M
V
M
V
M
V
M
V
M
V
M
V
-
Tn-RpL32 (260 bp)
CrPV_pair-5 (223 bp)
300 bp
200 bp
100 bp
1500 bp
1000 bp
850 bp
650 bp
500 bp
400 bp
300 bp
200 bp
100 bp
Figure S2: Presence of intracellular vDNA from CrPV in High Five cells upon infection. A: Amplicons produced with CrPV-specific primers (CrPV_pair-5) from DNA, RNA, or digested DNA, collected immediately (T0) or three days (T3) after viral infection (V) or mock-infection (M) of cells. B: Amplicons produced with TnRpL32-specific primers from DNA of cells collected immediately (T0) or three days (T3) after viral infection (V) or mock infection (M). -: non-template control. The Thermo Fisher Scientific 1 Kb Plus DNA ladder was used as a reference.

### Slide 4
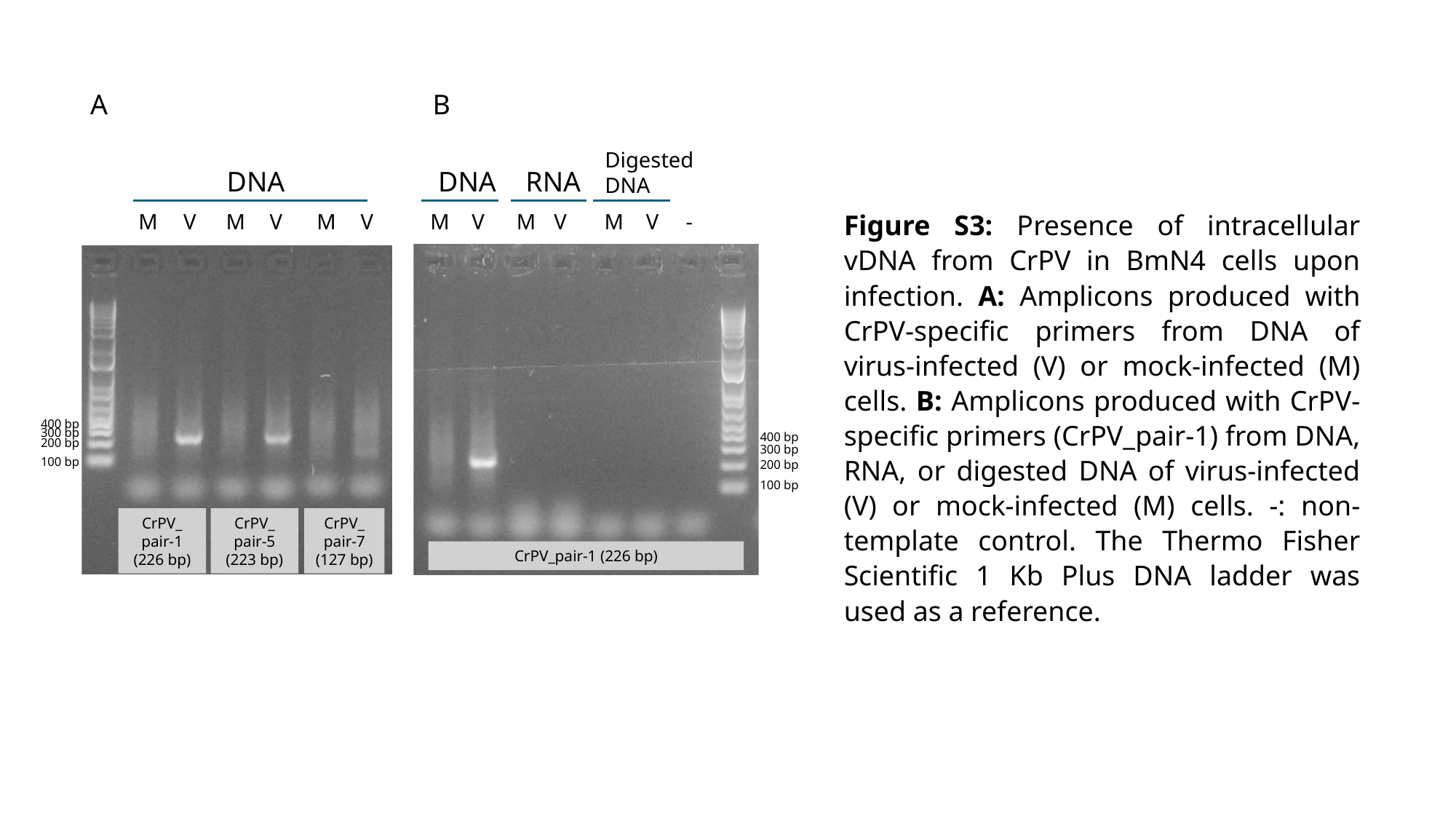

A
B
Digested DNA
DNA
DNA
RNA
Figure S3: Presence of intracellular vDNA from CrPV in BmN4 cells upon infection. A: Amplicons produced with CrPV-specific primers from DNA of virus-infected (V) or mock-infected (M) cells. B: Amplicons produced with CrPV-specific primers (CrPV_pair-1) from DNA, RNA, or digested DNA of virus-infected (V) or mock-infected (M) cells. -: non-template control. The Thermo Fisher Scientific 1 Kb Plus DNA ladder was used as a reference.
M
V
M
V
M
V
M
V
M
V
M
V
-
CrPV_
pair-1 (226 bp)
CrPV_
pair-5 (223 bp)
CrPV_
pair-7 (127 bp)
CrPV_pair-1 (226 bp)
400 bp
300 bp
400 bp
200 bp
300 bp
100 bp
200 bp
100 bp

### Slide 5
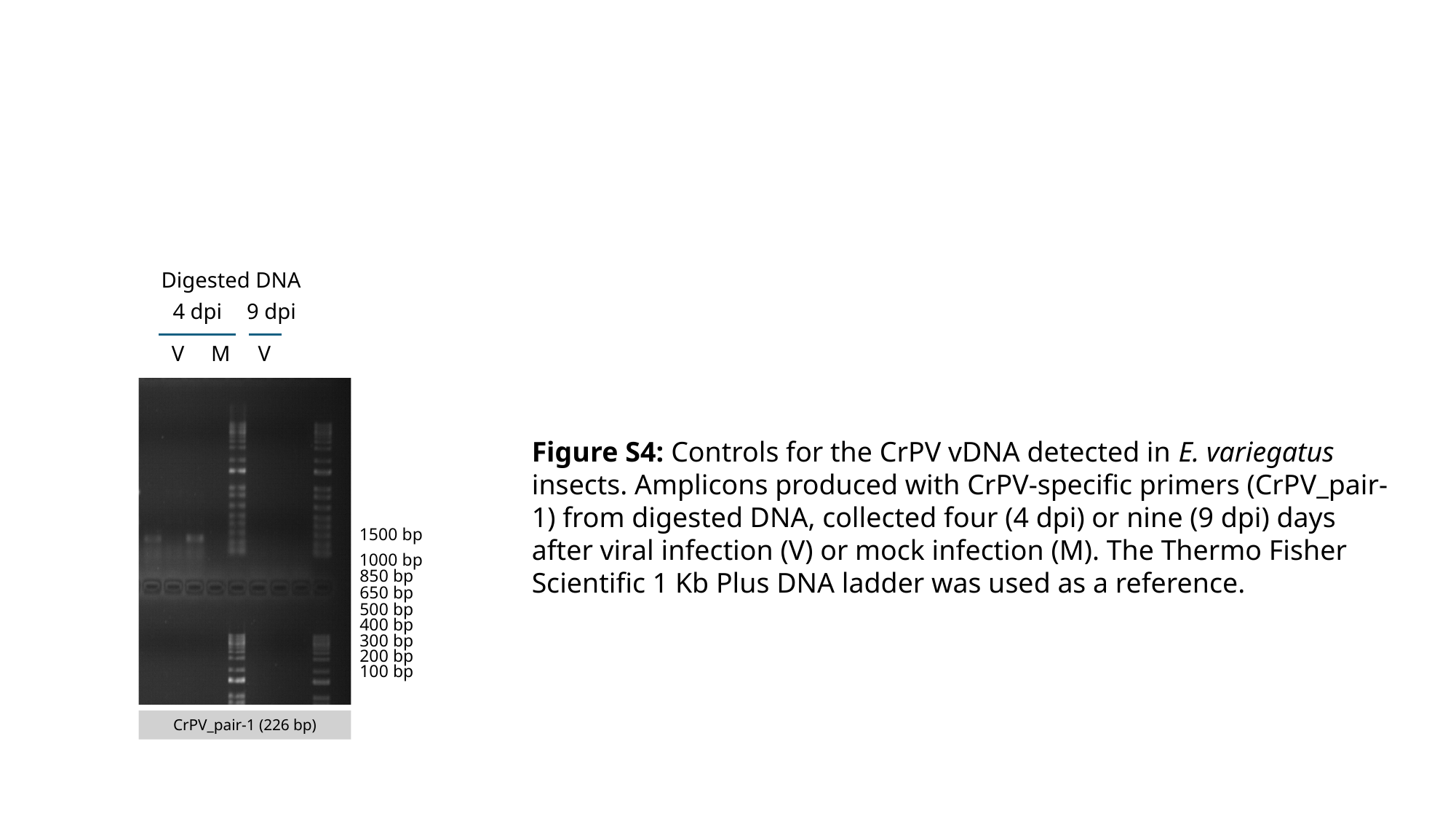

Digested DNA
4 dpi
9 dpi
V
M
V
Figure S4: Controls for the CrPV vDNA detected in E. variegatus insects. Amplicons produced with CrPV-specific primers (CrPV_pair-1) from digested DNA, collected four (4 dpi) or nine (9 dpi) days after viral infection (V) or mock infection (M). The Thermo Fisher Scientific 1 Kb Plus DNA ladder was used as a reference.
1500 bp
1000 bp
850 bp
650 bp
500 bp
400 bp
300 bp
200 bp
100 bp
CrPV_pair-1 (226 bp)

### Slide 6
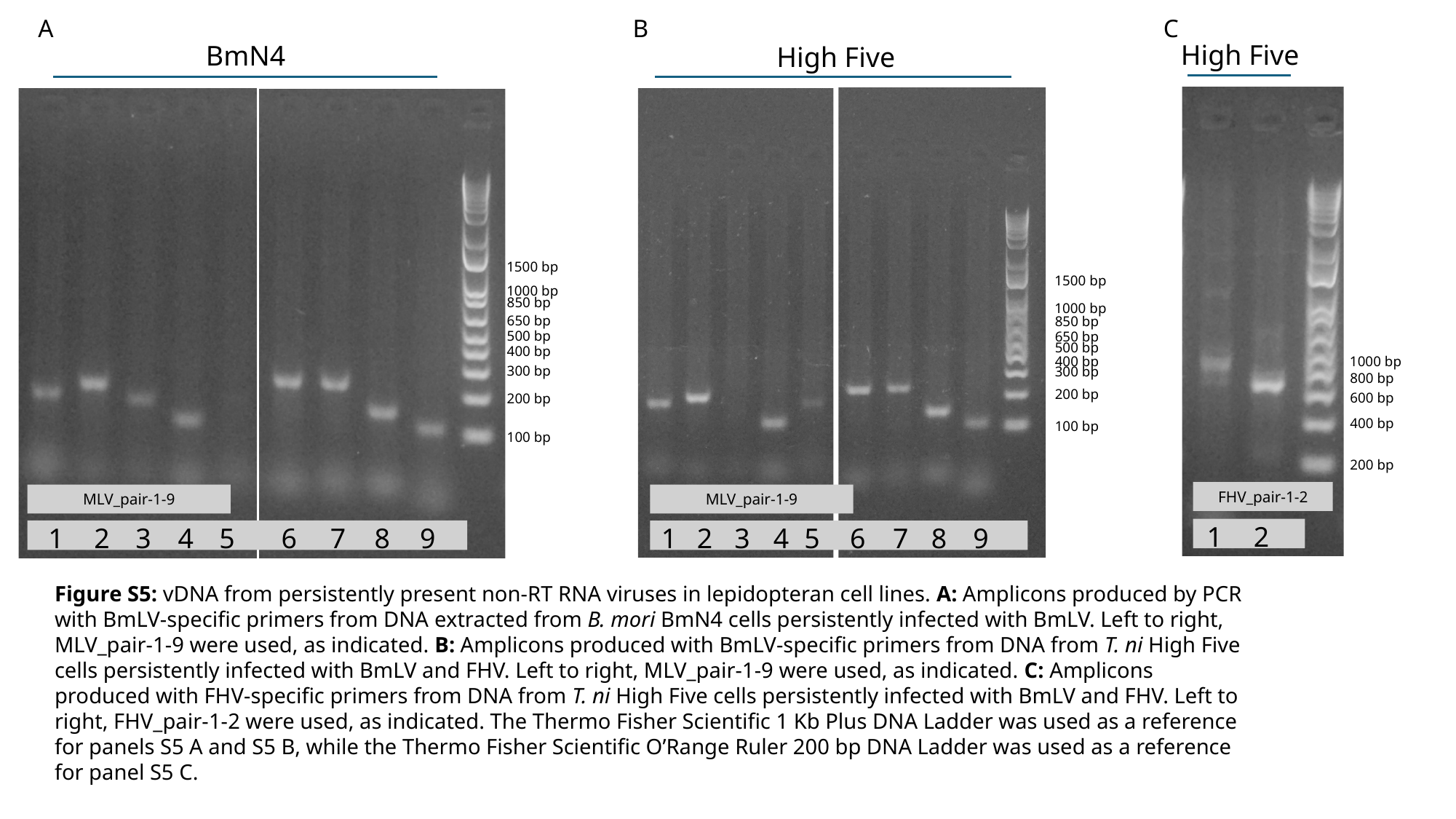

B
C
A
High Five
BmN4
High Five
1500 bp
1500 bp
1000 bp
850 bp
1000 bp
650 bp
850 bp
500 bp
650 bp
500 bp
400 bp
400 bp
1000 bp
300 bp
300 bp
800 bp
200 bp
600 bp
200 bp
400 bp
100 bp
100 bp
200 bp
FHV_pair-1-2
MLV_pair-1-9
MLV_pair-1-9
1
2
1
2
3
4
5
6
7
8
9
1
2
3
4
5
6
7
8
9
Figure S5: vDNA from persistently present non-RT RNA viruses in lepidopteran cell lines. A: Amplicons produced by PCR with BmLV-specific primers from DNA extracted from B. mori BmN4 cells persistently infected with BmLV. Left to right, MLV_pair-1-9 were used, as indicated. B: Amplicons produced with BmLV-specific primers from DNA from T. ni High Five cells persistently infected with BmLV and FHV. Left to right, MLV_pair-1-9 were used, as indicated. C: Amplicons produced with FHV-specific primers from DNA from T. ni High Five cells persistently infected with BmLV and FHV. Left to right, FHV_pair-1-2 were used, as indicated. The Thermo Fisher Scientific 1 Kb Plus DNA Ladder was used as a reference for panels S5 A and S5 B, while the Thermo Fisher Scientific O’Range Ruler 200 bp DNA Ladder was used as a reference for panel S5 C.

### Slide 7
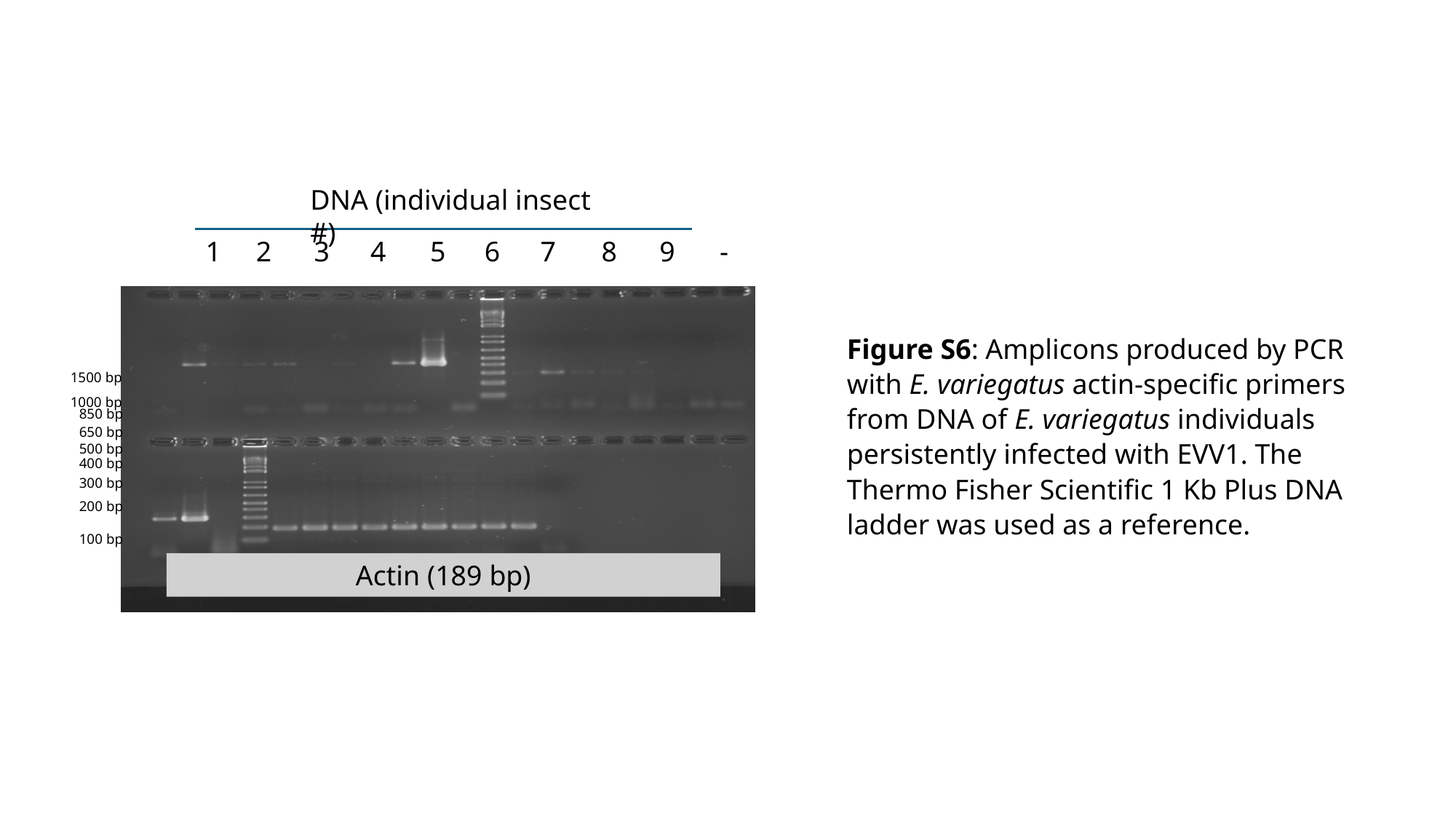

DNA (individual insect #)
1
2
3
4
5
6
7
8
9
-
Actin (189 bp)
Figure S6: Amplicons produced by PCR with E. variegatus actin-specific primers from DNA of E. variegatus individuals persistently infected with EVV1. The Thermo Fisher Scientific 1 Kb Plus DNA ladder was used as a reference.
1500 bp
1000 bp
850 bp
650 bp
500 bp
400 bp
300 bp
200 bp
100 bp
